# Luminance-corrected Pupillometry Reliably Tracks Effort in Dynamic Driving Environments

**DOI:** 10.64898/2026.05.29.728653

**Authors:** Yuqing Cai, Marnix Naber, Stefan Van der Stigchel, Christoph Strauch

## Abstract

Pupillometry provides an objective way to index effort across domains. However, pupil size is strongly affected by luminance changes, which can obscure effort-related effects, and limit its use in most applied scenarios with dynamic visual input. We here introduce and validate a method to overcome this problem. To this end, participants performed an auditory n-back task of differing difficulty while viewing either constant visual input or dynamic driving movie clips. Effort was assessed physiologically (pupil size), behaviorally (accuracy), and subjectively (NASA-TLX). Accuracy, questionnaire scores, and pupil size were analyzed at the session level, while pupillometry additionally provided continuous time-resolved information. As expected, pupillometry tracked differences in effort, but its discriminability was substantially reduced under dynamic visual input. Correcting for the effects of overall luminance and moment-to-moment luminance changes using a dynamic, explainable, and open-source modeling procedure (Open-DPSM) considerably improved effort discriminability on both aggregate and time-resolved levels. At the aggregate level, luminance-corrected pupillometry slightly outperformed accuracy and NASA-TLX. Combining all three measures yielded the highest classification performance (AUC = 0.98), supporting the view that effort is multifaceted and best captured multimodally. These findings establish a practical basis for fine-grained physiological tracking of effort and arousal in both fundamental and applied research using complex, dynamic stimuli. A tutorial section guides researchers in applying luminance correction to their own pupillometric data in dynamic viewing environments using the here validated approach.

## Introduction

Cognitive effort refers to the active allocation and deployment of limited cognitive resources, such as attention, working memory and executive control, to meet task demands, solve problems, or make decisions (Kahneman, 1973). Accurate measurement of effort is therefore essential for assessing task demands and understanding participants’ states, in fundamental research and applied fields alike. For instance, effort estimates may be used to inform adaptive systems across many domains, including driving, aviation, video gaming, education, and human–computer interaction more broadly, to enable real-time adjustments to task difficulty, feedback, or assistance based on the user’s current cognitive state (Dornhege et al., 2007; Evans & Fendley, 2017; Gabaude et al., 2012; Kantowitz & Casper, 2009; Kosch et al., 2023; Sevcenko et al., 2021). Effort is a multifaceted construct, composed of three main components: subjective experience (how hard does a task feel?), behavior (how is task performance affected?), and physiology (how does the body respond?) (Longo et al., 2022; Westbrook & Braver, 2015). Correspondingly, common measures of effort include subjective self-reports (e.g., NASA-TLX, Hart & Staveland, 1988)), behavioral performance (such as accuracy) and physiological measures (such as pupillometry or EEG) (O’Donnell & Eggemeier, 1986).

From a measurement perspective, physiological measures are perhaps the most challenging. This is not only due to the extra hardware and software requirements, but foremost due to the inherently complex and indirect nature of physiological signals. At the same time, physiological measures are in many contexts the most informative and promising, as they do not require explicit responses, can be assessed and analyzed continuously, and are not bound by subjective biases or limits to interoception (Diarra et al., 2025; Tao et al., 2019; van der Wel & van Steenbergen, 2018). A variety of physiological indices such as EEG, heart rate variability, skin conductance, and eye-tracking metrics, including pupil size change have all shown to reflect effort (Das Chakladar & Roy, 2024; Tao et al., 2019). Among these metrics, pupillometry is perhaps the most established: pupil size increases with any increase in effort (Beatty, 1982; Bumke, 1911), reliably found across various tasks and paradigms, such as mental arithmetic (Ahern & Beatty, 1979; Hess & Polt, 1964; Klingner et al., 2011), working memory (Kahneman & Beatty, 1966; Koevoet et al., 2024; Unsworth & Robison, 2015), and language comprehension (Just & Carpenter, 1993). Moreover, pupil sizes are measured remotely and unnoticeably, thus creating minimal interference to natural behavior. Effectively, this makes pupillometry particularly suitable for assessing effort in realistic, dynamic settings, in theory (Duchowski, 2017; Kosch et al., 2023; Skaramagkas et al., 2023).

Unfortunately, while the pupil has been found highly sensitive to even the finest changes in effort in the lab, at constant fixation and luminance (e.g., de Gee et al., 2014; Koevoet et al., 2025), it is a fundamentally integrated measure. This means that the pupil does not only respond to changes in effort, but foremost to brightness changes (see Mathôt & Van der Stigchel, 2015; Strauch et al., 2022; Woodhouse & Campbell, 1975 for reviews). Hereby, pupils constrict as luminance increases and dilate as luminance decreases. This pupillary light response is often much larger than cognitively driven changes in pupil size (Beatty, 1982; Mathôt, 2018). Hence, controlling luminance is the essential standard for pupillometric studies into cognitive processes in controlled lab settings (Beatty & Lucero-Wagoner, 2000; Mathôt & Vilotijević, 2023). Consequently, much fewer studies use pupillometry in unconstrained environments with dynamic visual input and few considered confounding effect of luminance. Little surprisingly, uncontrolled luminance leads to weak or inconsistent relationships between pupil size and effort (Evans & Fendley, 2017; Johannessen et al., 2020; Lin et al., 2008; van der Wel & van Steenbergen, 2018), unless being corrected for (Lee et al., 2024; Mitre-Hernandez et al., 2021; Wong et al., 2020). For many fundamental use-cases and most applications of effort measurements, a flexible and reliable pupil-luminance correction is thus of utmost importance if researchers want to study scenarios that go beyond looking at a centrally positioned dot with constant visual input.

One solution to this problem lies in an originally proprietary method (i.e., the Index of Cognitive Activity; Marshall, 2002, 2007) that now has an open-source counterpart (i.e., the Index of Pupillary Activity; Duchowski et al., 2018, 2020). These approaches try to obtain effort-related effects by decomposing the pupillary signal in the frequency domain (Duchowski et al., 2018). However, although they have shown sensitivity to effort in certain tasks (Devos et al., 2020; Fairclough et al., 2009; Schwalm, 2009), their generalizability across tasks, and validity under dynamic visual input, remain debated (Kahya et al., 2018; Korbach et al., 2017; Rerhaye et al., 2019; Schierhorst et al., 2024; Vogels et al., 2018).

More promising solutions may thus lie in accounting for the influence of luminance on the pupillary signal under dynamic visual input to remove luminance-induced effects and obtain clearer effort-related influences. One widely adopted and easy approach is to baseline correct, i.e., measuring pupil size during a baseline period and subtracting that size from the following data points recorded during a period the effect of effort is expected. This lab standard (see e.g., Mathôt, 2018) has also been adopted in studies with dynamic visual stimuli to reduce effects of individual differences and luminance-induced confounding (Kiefer et al., 2016; Liao et al., 2025; Mitre-Hernandez et al., 2021). However, its effectiveness in enhancing effort-related pupil signals has rarely been tested explicitly. Moreover, baseline correction can only control for the overall luminance level, but not for the moment-to-moment changes in luminance that characterize the ever-changing input (due to changes in the environment and eye movements). Effectively, this strips pupil size changes of one of its core advantages as a measure of effort, namely its often-highlighted continuous nature (Beatty, 1982; Granholm & Steinhauer, 2004; van der Wel & van Steenbergen, 2018; Wong et al., 2020).

A much smaller number of studies modeled pupil size changes to moment-to-moment luminance changes in the visual input to regress out luminance-related changes, yielding a ‘purified’ continuous effort signal. For example, previous driving studies illustrate both the need and the feasibility of decoupling luminance-driven from effort-related pupil responses by modeling the pupillary light reflex (Kun et al., 2012; Palinko & Kun, 2011). However, these studies rely on simple stimuli (static images or constrained visual scenes) rather than dynamic visual input and did not quantify whether the luminance correction actually improved effort estimation. More recent studies have extended luminance modeling to dynamic stimuli (Fanourakis & Chanel, 2022; Napieralski & Rynkiewicz, 2019; Raiturkar et al., 2016). Those models can be categorized into either steady-state models, which predict pupil size from luminance in short discrete intervals (Chen et al., 2017; Fanourakis & Chanel, 2022; John et al., 2018; Napieralski & Rynkiewicz, 2019; Raiturkar et al., 2016; Watson & Yellott, 2012), and dynamic models, which model temporal dynamics of pupil responses to transient luminance changes (Cai et al., 2024; Fan & Yao, 2011; Fanourakis & Chanel, 2022; Pamplona et al., 2009). Notably, only few studies quantified whether removing luminance confounds actually improved effort detection (Fanourakis & Chanel, 2022; Raiturkar et al., 2016). Moreover, for dynamic visual input, dynamic models are naturally more effective, but adoption remains low (Fanourakis & Chanel, 2022). This is partly because many of these models are based on biomechanical functions of the eye and brain and thus are mathematically complex (with many parameters to estimate) and difficult to implement due to a lack of accessible toolboxes (Soleymani et al., 2012; Zandi & Khanh, 2021).

An alternative lies in convolutional models. Convolutional models take luminance changes and multiply them with the known shape of pupil responses to luminance changes and aggregate this over time (i.e., a response function; Hoeks & Levelt, 1993; Korn & Bach, 2016). Recently, Cai et al. (2024) put forward such a model, coming with completely open-source code, a toolbox, and an accessible graphical user interface (Open-DPSM; https://github.com/caiyuqing/Open-DPSM). Subsequent studies have also shown that this approach can, for instance, improve indices of visual sensitivity and covert attention under dynamic visual input (Cai, Strauch, et al., 2025; Cai, Van der Stigchel, et al., 2025). Open-DPSM requires nothing but a video of the visual input along with gaze positions and pupil size changes obtained by an eye-tracker. Whether Open-DPSM can improve the measurement of effort under dynamic visual input will be directly tested in the present paper. Most recently, first evidence emerged from an independent lab, demonstrating that Open-DPSM improves arousal rating prediction when watching videos (Wang et al., 2026).

The current study examined pupil size as an index of cognitive effort under dynamic and highly complex visual input. Rather than predicting arousal with pupil size (Wang et al., 2026), our work directly aims at correcting for low-level pupillary effects to better capture mental effort. Furthermore, we provide fully open code and an extension built on the most recent version of Open-DPSM. Specifically, we chose driving because pupillometry has been widely used to assess effort in this field or its use been called for (Beggiato et al., 2018; J. Huang et al., 2024; Palinko et al., 2010; Prabhakar et al., 2020; Radhakrishnan et al., 2022, 2023; Rendon-Velez et al., 2016; Vintila et al., 2017), but note that results will similarly apply to *any* dynamic visual input, whether it is a video intended to induce emotional arousal, educational material, or a video game. To this end, we manipulated effort using an auditory n-back task with either constant visual (mimicking classic lab studies as a baseline) or dynamic driving videos with a high degree of changes in visual input (mountain roads, forests, cities etc.) to test the method with the hardest possible visual input. We tested whether luminance correction would improve pupil sensitivity to effort at two temporal scales: (1) on a session level (low temporal resolution), overall luminance correction was applied to the aggregated pupil size; and (2) over time (high temporal resolution), using Open-DPSM to estimate and remove luminance-induced pupil size changes. To capture effort fully and as a benchmark to pupillometry, subjective and behavioural measures were also included. Finally, we tested whether a combination of measures would provide a superior estimation of effort, as to be expected from its multidimensional nature (Dhengre & Rothrock, 2025; J. Huang et al., 2024; Mitre-Hernandez et al., 2021).

## 2. Methods

### 2.1 Participants

36 healthy students (8 men; 28 women, *M_age_* = 22.83, *SD* = 5.49) with normal or corrected-to-normal vision participated in the experiment after providing informed consent. Two participants were removed due to excessive missing pupil data (one due to keep looking down at the keyboard and the other due to unsuccessful calibrations; 56.75% and 47.86%), resulting in a final sample of 34 participants. This study was approved by the local ethical committee of the authors’ institution (#22-530).

### 2.2 Apparatus

The experiment was conducted in a laboratory with controlled ambient lighting (the luminance of the wall behind the monitor was 2.57 cd/m²). The task was programmed in PsychoPy (version 2024.2.4; Peirce et al., 2019) and presented on a 34.5 x 59.5cm (32.08° by 52.74° visual angle, 1920 x 1080 pixels) monitor with a refresh rate of 60 Hz. Eye movements and pupil diameters were recorded using a Tobii Pro Spectrum eye-tracker at 300 Hz. Participants were seated approximately 60 cm from the monitor with a chin rest. Gaze position was calibrated using a 9-point calibration and validation procedure with the Python Titta package (Niehorster et al., 2020).

### 2.3 Stimuli and procedure

Participants performed an auditory n-back task with three levels of difficulty to manipulate their memory load (0-back, 1-back, and 2-back, see Huang et al., 2025; Owen et al., 2005 for reviews on the n-back task). Participants heard a series of spoken digits (0-9) in random order, presented in English by a synthetic female voice. The volume was individually adjusted to a comfortable level for each participant. Each digit was played for 1 s, followed by a 2 s interstimulus interval. In the 1-back and 2-back conditions, participants were instructed to press the space bar whenever the current digit matched the digit presented one or two trials earlier, respectively. For the 0-back condition, a predefined target digit was randomly assigned before each session and the participant responded whenever they heard that digit. Each session contained 100 digits and target digits constituted 20% of all the trials for each session, in random temporal sequence.

To provide a controlled laboratory benchmark, participants performed one session per n-back condition with constant gaze and visual input (a black screen (0.87 cd/m2, measured at the participant’s eye position)) with a white central fixation cross (47.3 cd/m^2^, measured at the participant’s eye position; constant input condition, see Fig. 1 for an illustration of the experimental paradigm). Participants were instructed to keep looking at the center cross. To put pupillometry to a hard test with dynamic visual input, participants watched driving movie clips while performing the n-back task for six sessions, with two sessions for each n-back condition (dynamic input condition; see Fig. 1). Participants were instructed to simply view the movie while performing the auditory task. Six driving movie clips (5 minutes each, CC-BY license) were downloaded from YouTube. The clips depicted recordings of car drives, filmed from the front of the vehicle, driving through different, complex environments, such as a city, a mountain road, etc. (see https://osf.io/2tbpk component “Driving videos” for movies).

**Figure 1.**
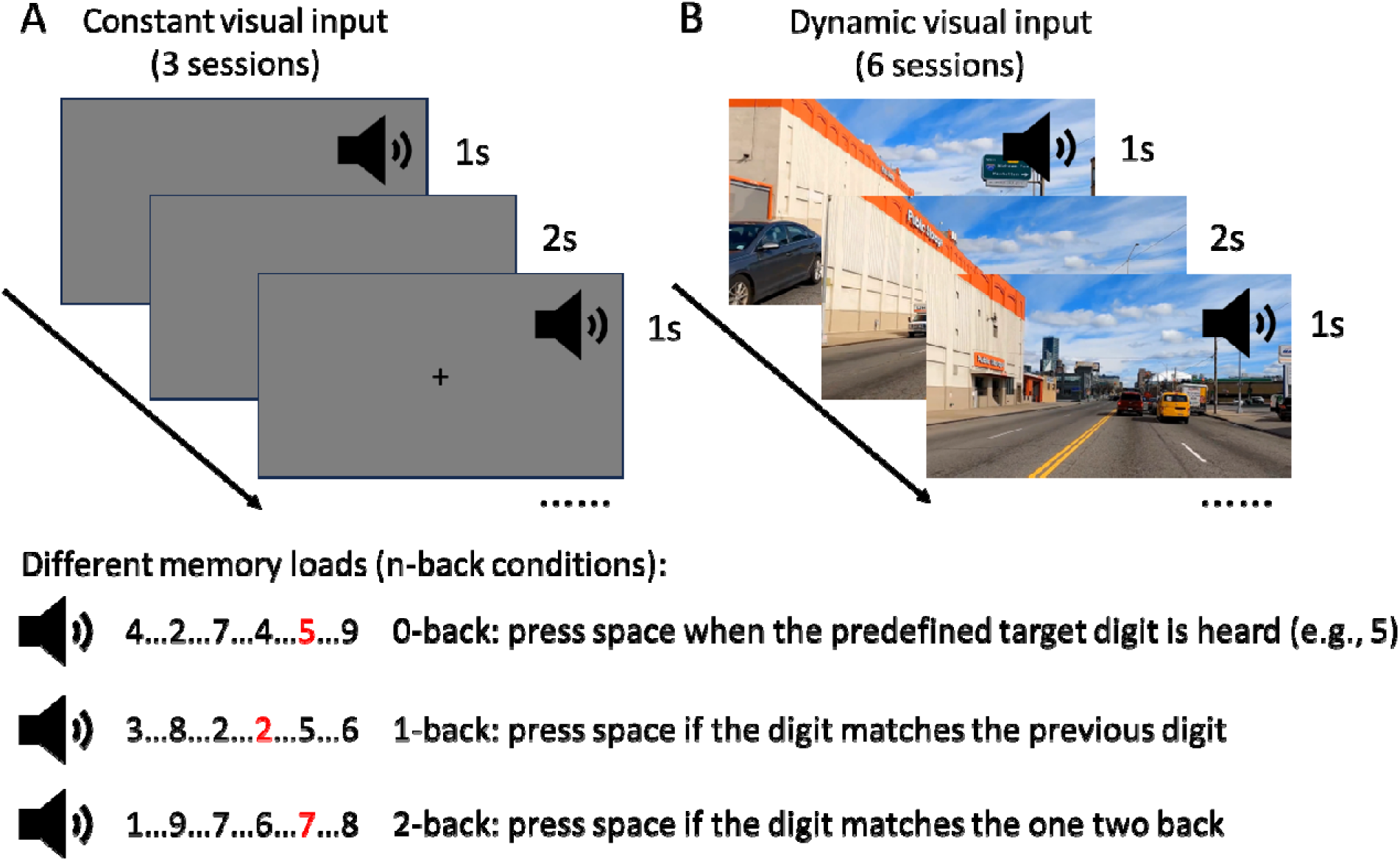
Experimental paradigm. *Note: Participants performed a n-back task ofthree different loads while being presented with constant visual input (A) or with dynamic visual input in the form ofa driving movie clip (B). Texts at the bottom indicate tasks across n-back conditions, with red indicating exemplary target digits.*

Participants first completed three short practice sessions (20 digits each), one for each n-back condition without the movie to familiarize themselves with the task. During the main experiment, the order of conditions and movie clips assigned were fully randomized across participants.

After each session, participants completed the NASA Task Load Index (NASA-TLX; Hart & Staveland, 1988), a standard self-report measure of subjective workload, with six sub-scales assessing mental demands, physical demands, temporal demands, own performance, effort, and frustration. Participants scored each item with a visual analogue scale ranging from 0 to 100 in 5-point increments on screen. Note that the original questionnaire also contained a weighing procedure to determine the perceived importance of each sub-scale, but in this study, the unweighted raw NASA-TLX score was used because previous studies have shown that the unweighted average provides a simple and sensitive measure of workload, with high test-retest reliability (Devos et al., 2020; Ikuma et al., 2009; Said et al., 2020).

### 2.4 Data analysis

Measures of cognitive effort were divided into two categories based on their temporal resolution. Aggregate measures that provided a single measurement value per session were considered as low in temporal resolution, including accuracy, NASA-TLX scores, and mean pupil size per session. Time-series pupil size provided a measurement for each time point and was thus used as a high-temporal resolution measure, potentially allowing moment-to-moment assessment of cognitive effort.

How well those measures reflect (differences in) cognitive effort across n-back conditions was assessed. Furthermore, discriminability analyses were performed to determine whether, and if so, how well, those measures can distinguish different n-back conditions. To evaluate the influence of luminance, we compared the results of constant input and dynamic input conditions, and assessed whether luminance correction would allow the inference of effort from pupil size under dynamic visual input. For mean pupil size, overall luminance correction was applied for each session. For time-series pupil size, we applied dynamic luminance correction using a recent, open-source, and state-of-the art model (Cai et al., 2024).

## Results

### 3.1 Preprocessing

One session was excluded due to hardware issues, resulting in 305 sessions in total from 34 participants.

Blinks were identified with the eye-openness parameter provided by Tobii, as well as pupil size change velocities (values exceeding three standard deviations from the mean velocity were classified as blinks). Blink periods were removed and interpolated. The data were then down-sampled to 25 Hz, to match the movie frame rate (25 fps). Sessions with more than 40% of missing data after blink removal (before interpolation) were excluded (8.85% of sessions), resulting in 278 usable sessions.

### 3.2 Aggregate measures of cognitive effort

#### 3.2.1 Accuracy and NASA-TLX

Both accuracy on the n-back task and NASA-TLX provided (only) low-temporal resolution aggregate per-session measures. Accuracy was defined as the proportion of correctly detected target trials (number of correctly identified targets / total number of targets). Sessions with accuracy below 0.5 were removed (1.07% of sessions).

Accuracy and NASA-TLX were analyzed using a repeated-measures ANOVA with visual input conditions (constant input vs. dynamic input) and n-back level (0-, 1-, and 2-back) as within-subject factors and post-hoc comparisons were made with paired-sample t-tests. Multiple comparisons were corrected with Holm method and effect size was quantified with Cohen’s d. Only participants with full data for each condition were analyzed. Sphericity was evaluated with Mauchly’s test of sphericity for all the ANOVA analyses with factors containing more than two levels, and none of the ANOVAs reported below violated sphericity.

Repeated measures ANOVAs revealed significant main effects of n-back load on both accuracy and NASA-TLX (both *p* < 0.001) with no main effects for visual input (both *p* > 0.06) and no interactions of the two (both *p* > 0.67; see Supplementary Table 1 for full statistics). To verify that the n-back manipulation successfully varied cognitive effort, we compared accuracy and NASA-TLX scores across n-back conditions. As expected, accuracy was lower in the 2-back condition, compared with 1- and 0-back conditions, but was not different between 0- and 1-back conditions (Fig. 2A-B; see Supplementary Table 2A for full statistics). NASA-TLX score also showed that the 2-back condition was perceived as more difficult than 1-back/0-back conditions. In addition, 1-back was also perceived as more difficult than 0-back in the dynamic input condition (Fig. 2C-D; see Supplementary Table 2B for full statistics).

**Figure 2.**
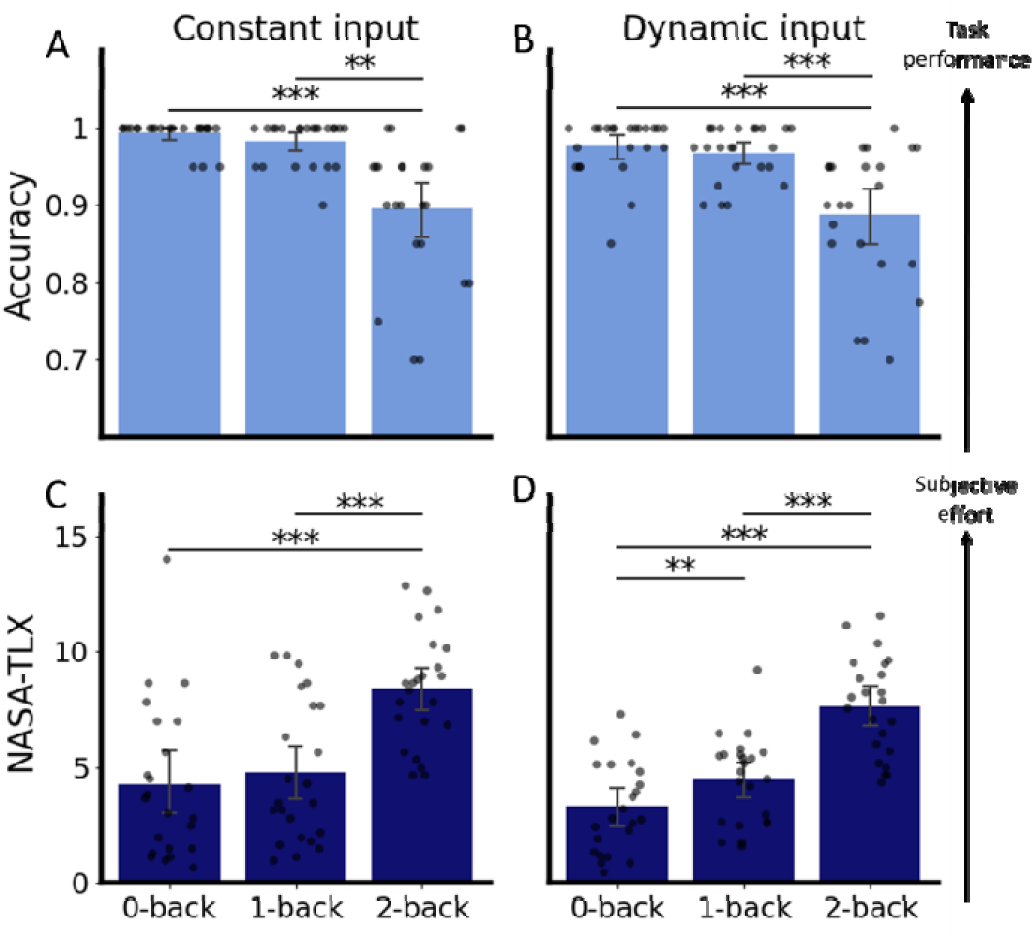
Accuracy and NASA-TLX score as measurements of cognitive effort. *Note: Each dot is one participant. Error bars represent 95% confidence intervals ofthe mean. ***: p < 0. 001, **: p < 0. 01*

Overall, accuracy and NASA-TLX score confirmed that the manipulation of cognitive effort was successful: 2-back condition showed lower accuracy and higher perceived effort ratings than the 1- and 0-back conditions. In addition, the 1-back condition was perceived as more difficult than the 0-back condition, but only under dynamic visual input.

#### 3.2.2 Mean pupil size

Mean pupil size was computed as the average pupil size after excluding the first 9s of each session (corresponding to the first three trials, during which cognitive effort was not yet fully manipulated through the n-back conditions). We first corrected for individual differences by subtracting the mean pupil size per participant (following e.g., Kun et al., 2012; Palinko & Kun, 2011; Pfleging et al., 2016).

To mitigate the effects of luminance, a linear mixed-effects model was used to predict mean pupil diameter with time-averaged overall luminance of the movie, with participant as a random intercept:

*Mean pupil size ∼ overall luminance (per movie) + (1|participant)*

This overall luminance model was used to capture session-level differences in pupil size, induced by different overall luminance levels across movies. The predicted pupil size of each session was then subtracted to obtain a luminance-corrected measure (a similar regression approach can be found in Hansen et al., 2018).

One-way repeated-measures ANOVAs revealed significant main effects of n-back load on mean pupil size, for constant input, dynamic input, and dynamic input with overall luminance correction (all *p* < 0.001; see Supplementary Table 3 for full statistics). To examine whether pupil size can index effort under dynamic input, we compared pupil size across n-back conditions. Consistent with results of accuracy and NASA-TLX, for both constant input and dynamic input, pupil size was larger in the 2-back condition than 1-back/0-back conditions (all *p* < 0.001), but was not different between 1-back and 0- back conditions (both *p* > 0.26), indicating that pupil size reflected effort, also when visual input was complex and dynamic (Fig. 3A-B; see Supplementary Table 4 for full statistics).

**Figure 3.**
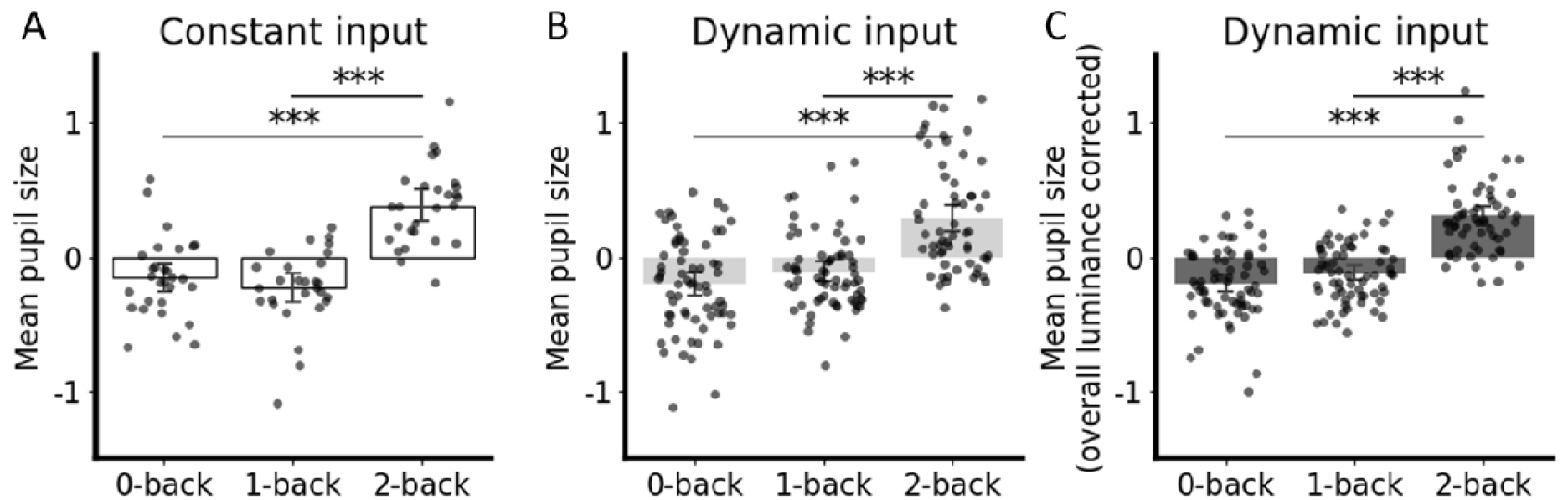
Mean pupil size as aggregate measure of cognitive effort. *Note: All pupil size measurements were first corrected for individual differences by subtracting average pupil size per participant. Each dot is one participant. Error bars represent 95% confidence intervals of the mean. ***: p < 0. 001*

After correcting for overall luminance, pupil size remained larger in the 2-back condition, compared with both 1- and 0-back conditions (both *p* < 0.001; Fig. 3C; see Supplementary Table 4 for full statistics). Crucially so, the effect size (*d*) was much larger than that for pupil size without luminance correction (uncorrected: d = -0.93 and -0.98, overall luminance-corrected: *d* = -1.37 and -1.38, for 2- vs. 1-back and 2- vs. 0-back respectively), suggesting that overall luminance correction reduced luminance-related confounding and enabled effort-related effects on pupil size easier to detect.

#### 3.2.3 Associations between pupillometric indices, and behavioral and subjective measures

Given the multifaceted nature of effort, the aforementioned measures should be linked. To explore such a relationship between pupillometric indices and behavioral/subjective indices, Pearson’s correlations were computed between pupillometric indices and accuracy as well as NASA-TLX across all sessions, pooling data from all n-back conditions (not on condition-averaged values). Mean pupil size was correlated with accuracy/NASA-TLX score (see Table 1 for details). Unsurprisingly, this correlation was strongest under constant visual input. For dynamic visual input, this correlation became higher when the pupil size was corrected for luminance. Accuracy was also correlated with NASA-TLX.

**Table 1.** Correlation (*r*) between pupil size, accuracy and NASA-TLX.

|  | Accuracy | NASA-TLX |
| --- | --- | --- |
| Pupil size (constant input) | -0.50*** | 0.42*** |
| Pupil size (dynamic input) | -0.31*** | 0.31*** |
| Pupil size (dynamic input, overall-luminance removed) | -0.33*** | 0.37*** |
| NASA-TLX | -0.51*** | - |

Note: \*\*\*: $p < 0.001$

#### 3.2.4 Mean pupil size discriminates n-back conditions better than accuracy and NASA-TLX

The preceding ANOVA and post-hoc tests demonstrated that 1-back and 0-back did not differ on most measures. Hence, subsequent analyses focus on pairwise comparisons between 1- and 2-back conditions, as well as 0- and 2-back conditions.

To quantify how well each measure discriminated between n-back conditions, univariate discriminability analyses were conducted within a signal-detection framework. Two signal detection metrics, area under the curve (*AUC*) of the receiver operating characteristics (ROC) and *d’*, the distance between the peaks of the distributions in z-scored indexes, were calculated. Analyses were performed at the session level, pooling across participants. For each measure, the decision threshold was selected by maximizing Youden’s *J* statistic.

Under constant visual input, accuracy, NASA-TLX, and pupil size all discriminated 2-back condition from the lower-load conditions (all *AUC* > 0.79, *d’* > 1.49; Fig. 4A-B; see Supplementary Table 5 for full statistics). Similarly, under dynamic visual input, accuracy, NASA-TLX score, and pupil size, as well as pupil size corrected for overall luminance all discriminated 2-back from 0-back and 1-back conditions (all *AUC* > 0.77, *d’* > 1.22; Fig. 4C-D; see Supplementary Table 5 for full statistics). Notably, mean pupil performed numerically even better than accuracy and NASA-TLX in the constant input condition. While this measure performed slightly worse compared to accuracy and NASA-TLX score in the dynamic input condition, simple luminance correction let it again numerically outperform accuracy and NASA-TLX score (see Fig. 4 for illustration and Supplementary Table 5 for full statistics).

**Figure 4.**
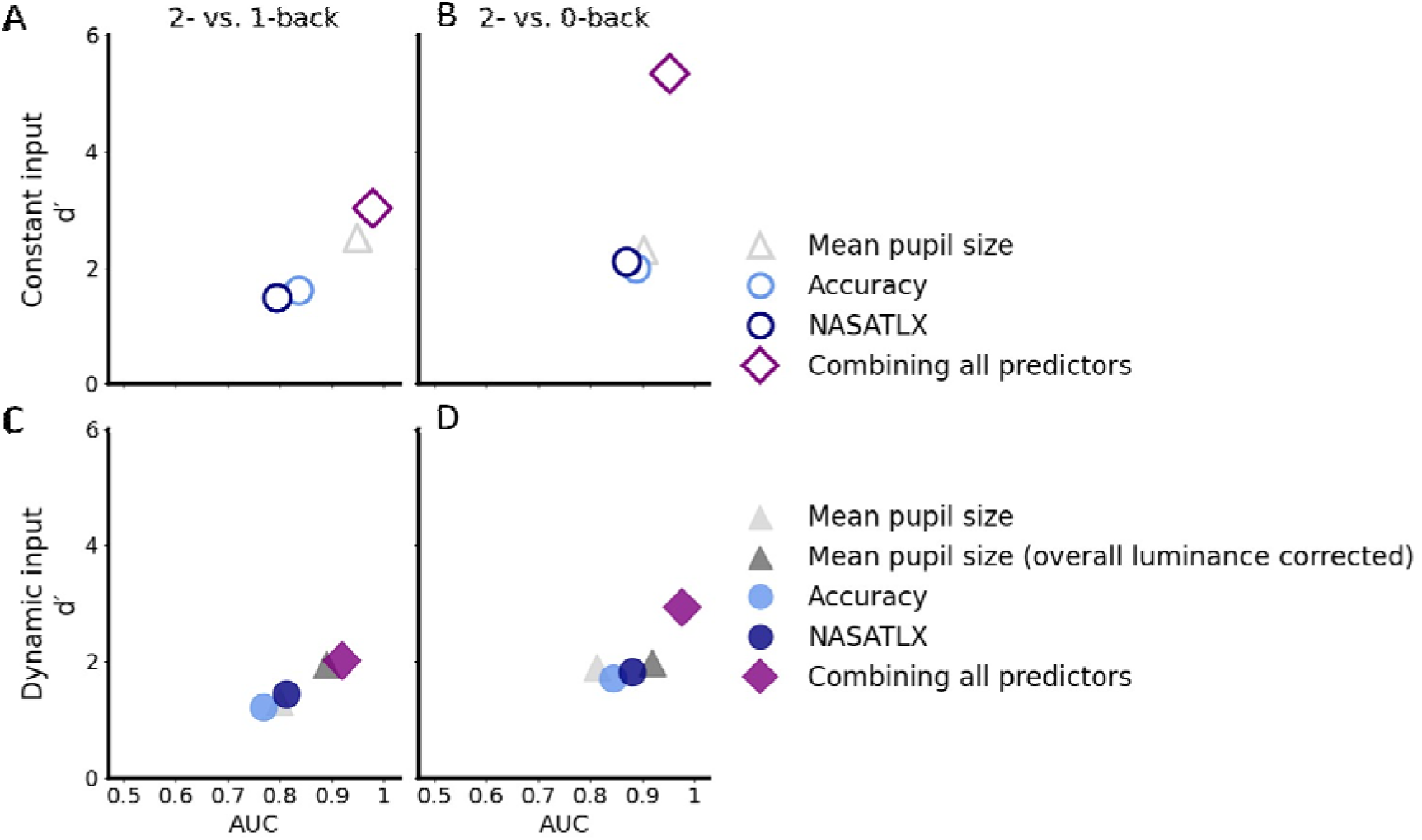
Discriminability of all measurements tested. *Note: The discrimination test was performed on all sessions, pooled across participants. Triangles: pupil size, circles: other factors. Diamond: all factors combined. Top right indicates the best discrimination performance*.

#### 3.2.5 Combining physiological, behavioral, and subjective indices provides superior effort measurement

Theoretically, a measure combining subjective, behavioural, and physiological aspects of effort should best dissociate differing degrees of task load/effort. To assess whether combining pupillometric, behavioural and subjective measures yielded a better classification result for effort, a multivariate classification analysis with logistic regression, with all factors as predictors, was conducted. Logistic regression classifiers were trained to distinguish 2- from 1-back or 0-back conditions. For the constant input condition, the predictors were mean pupil size, accuracy and NASA-TLX score and for the dynamic input condition, the predictors were mean luminance-corrected pupil size, accuracy, and NASA-TLX score. All predictors were standardized (z-scored) prior to model fitting.

Classification performance was evaluated using leave-one-subject-out cross-validation, such that sessions from one participant were excluded from model training and were used exclusively for testing. Performance (_AUC_ and d□) was based on pooled out-of-sample predictions across all cross-validation folds. Results illustrated that combining behavioural, subjective and pupillometric measures improved classification performance compared to any single measure alone for both constant and dynamic visual input conditions (all *AUC* > 0.92, *d’* > 2.02; see Fig. 4 for illustration and Supplementary Table 5 for full statistics), indicating that these measures partially captured different aspects of cognitive effort. Given that effort naturally varies in each session across participants (due to differing levels of task engagement, commitment, attention etc.), an AUC of 1.00 would be theoretically impossible to achieve. Thus, the near-perfect discrimination between conditions observed in our classification analyses was potentially already providing perfect effort separation.

#### 3.2.6 Pupillometry provides unique information about cognitive effort beyond behavioural and subjective measurements

We showed that combining pupil size, accuracy and NASA-TLX improved classification compared with using any single measure alone. However, it remains unclear whether pupil size captures distinct variance of cognitive effort, when others were controlled. Hence, we used logistic regression models with L2 (ridge) regularization to examine the unique contribution of pupil size after controlling for other two variables, and to quantify the relative importance of all three measures in explaining variance. The outcome variable (n-back condition) was dichotomized into high load (2-back) versus low load (0-/1-back). All predictors were standardized (z-scored) prior to model fitting.

Ridge regularization was adopted to mitigate the influence of multicollinearity among predictors and to obtain stable coefficient estimates. The strength of regularization (inverse penalty parameter C) was selected by minimizing cross-validated loss using 5-fold cross-validation.

Results demonstrated the unique contribution of mean pupil size under both constant input and dynamic input (Table 2 and Table 3), after accounting for other measures.

**Table 2.** Logistic ridge regression model for constant input condition.

| | Coefficient $\beta$ | Standard error | p-value | 95%CI |
| --- | --- | --- | --- | --- |
| Intercept | -1.63 | 0.52 | <0.001 | [-2.64, -0.62] |
| Accuracy | -1.25 | 0.50 | 0.01 | [-2.22, -0.28] |
| NASA-TLX | 0.85 | 0.44 | 0.05 | [-0.02, 1.72] |
| Pupil size | 2.36 | 0.67 | <0.001 | [1.05, 3.67] |

**Table 3.**
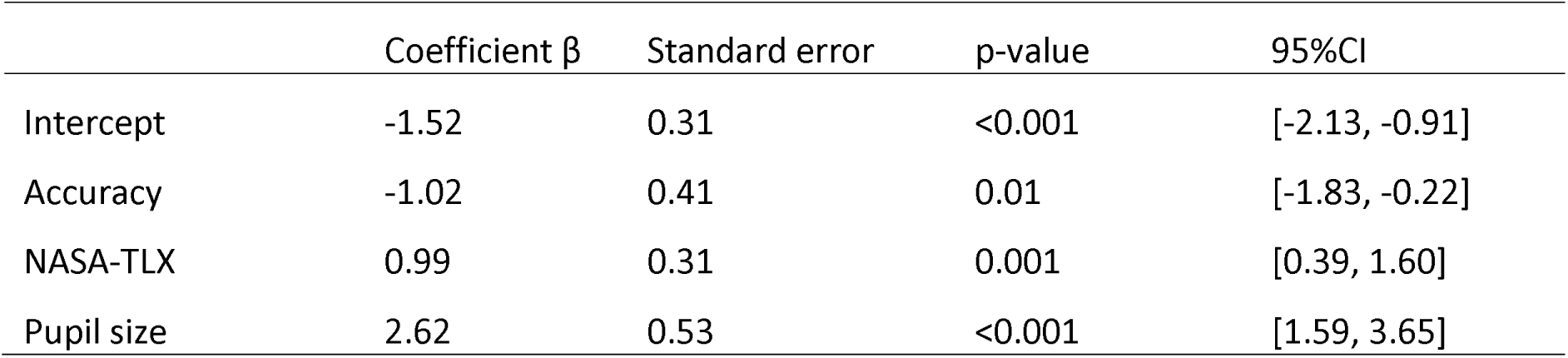
Logistic ridge regression model for dynamic input condition.

| | Coefficient $\beta$ | Standard error | p-value | 95%CI |
| --- | --- | --- | --- | --- |
| Intercept | -1.52 | 0.31 | <0.001 | [-2.13, -0.91] |
| Accuracy | -1.02 | 0.41 | 0.01 | [-1.83, -0.22] |
| NASA-TLX | 0.99 | 0.31 | 0.001 | [0.39, 1.60] |
| Pupil size | 2.62 | 0.53 | <0.001 | [1.59, 3.65] |

In addition, mean pupil size exhibited the largest standardized regression coefficient (Table 2 and Table 3), indicating that it was most strongly associated with cognitive effort relative to behavioral and subjective measures.

### 3.3 Estimating cognitive effort with pupil size over time

Our results, in line with previous works (Dhengre & Rothrock, 2025; J. Huang et al., 2024; Mitre-Hernandez et al., 2021), show that effort is multidimensional and is best characterized by combining measures of different modalities. However, when pupil size was used as an aggregate measure on session level, its phasic dilations and constrictions canceled out each other, making it impossible to capture fluctuations in effort. Yet, a decisive argument for why pupillometry is particularly promising in tracking effort is that pupil size provides a continuous, high-temporal resolution signal. To assess whether and how well pupil size could signal effort over time, univariate discrimination analyses were performed independently per time point (similar to that in section 3.2.4) (Fig.6A).

To correct for the influence of moment-to-moment luminance changes, a dynamic convolutional model (Open-DPSM; Cai et al., 2024) was applied. For the current study, we used an updated version of Open-DPSM. Shortly, the original Open-DPSM uses a response function (Fig. 5A) to approximate prototypical pupillary responses to luminance changes and convolves it with luminance changes extracted from the visual input (Fig. 5B). Luminance changes were extracted in a gaze-contingent manner, controlling for the influence of gaze position on luminance effectively reaching the retina. Hence, the model predicts the changes in pupil size driven by dynamic changes in luminance. However, correcting for luminance-driven effects on the observed pupil signal requires predicting absolute pupil size, rather than only pupil size changes. Therefore, in the updated version of Open-DPSM, we additionally included the predicted pupil size from the overall luminance model (section 3.2.2) as a known fixed component. The final model prediction therefore represents absolute pupil size, combining both time-averaged luminance effect and moment-to-moment dynamic luminance effects (Fig. 5C, dashed line). Subtracting the predicted luminance-driven pupil size from the observed pupil size (Fig. 5C, solid line) yielded a luminance-corrected pupil trace, which was used as a proxy for effort-related pupil activity (Fig. 5D).

**Figure 5.**
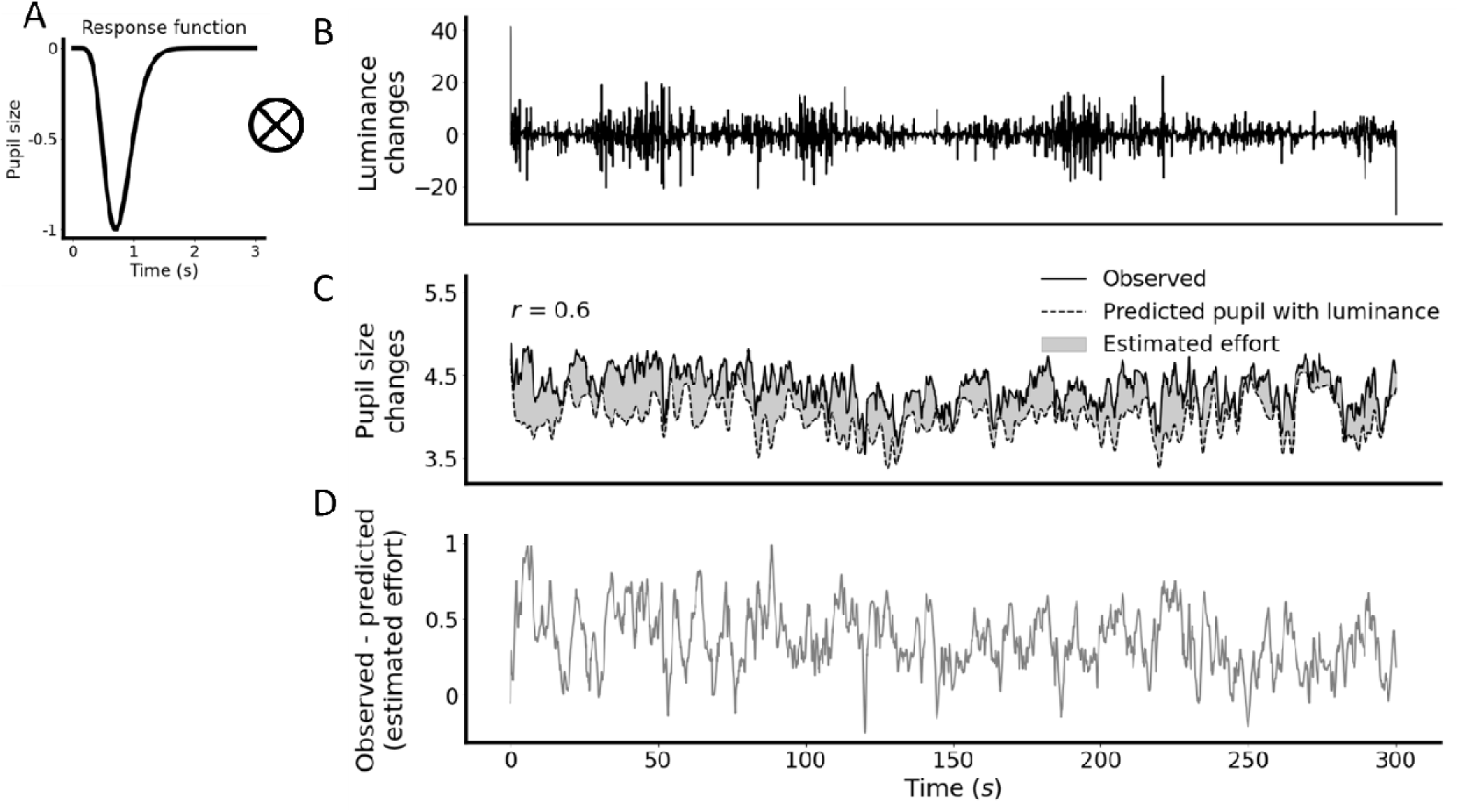
Illustration of how Open-DPSM removes luminance-change related effects on pupil diameter. *Note: Illustration of how Open-DPSM corrects pupil size for luminance-effects. Exemplary data from one session. A Prototypical pupillary response to luminance changes (response function); B Luminance changes extracted from the movie at each time point; C Observed pupil size changes (solid line) and pupil size predicted from luminance (dashed line) and their correlation (r = 0. 58). The predicted pupil size combined both the influence of overall luminance and the phasic influence of luminance changes. The shaded area represents the difference between observed and luminance-predicted pupil size; D Residual pupil size after removing luminance-driven components, corresponding to the shaded area in C, which can be interpreted as non-luminance-related pupil size change, predominantly reflecting task-related effort.*

Each session was modeled separately. The model was optimized by searching through different combinations of free parameters (e.g., parameters in the response function) to maximize the Pearson correlation between predicted and observed pupil size change. Correlation rather than root mean squared error (RMSE) was used as the optimization criterion to reduce the risk that sustained effort-related differences between n-back conditions would be absorbed into the luminance model. This updated version of toolbox is available at http://github.com/caiyuqing/Open-DPSM/tree/master/v4 (see next section for details).

Three approaches were compared to evaluate whether luminance correction would improve high-temporal resolution effort tracking under dynamic visual input: (1) pupil size without luminance correction; (2) pupil size corrected for the overall luminance, and (3) pupil size corrected with Open-DPSM, accounting for both overall luminance and luminance changes over time.

Time-resolved *AUC* and *d’* were computed for each time point first (Fig. 6A) and averaged across all time points (Fig. 6B). In the dynamic input condition, pupil size discriminated between n-back conditions (2- vs. 1-back: *AUC* = 0.71, *d’* = 1.04; 2- vs. 0-back: *AUC* = 0.74, *d’* = 1.19; grey triangles in Fig. 6B). Discriminability improved after correcting for overall luminance (2- vs. 1-back: *AUC* = 0.75, *d’* = 1.21; 2- vs. 0-back: *AUC* = 0.79, *d’* = 1.39; black triangles in Fig. 6B), and even further improved after applying Open-DPSM (2- vs. 1-back: *AUC* = 0.78, *d’* = 1.36; 2- vs. 0-back: *AUC* = 0.82, *d’* = 1.55; green triangles in Fig. 6B). Little surprisingly, the constant visual input condition yielded the numerical best discrimination, but note that values were quite close for fully-luminance corrected pupil size (constant visual input: 2-vs. 1-back: *AUC* = 0.81, *d’* = 1.67; 2- vs. 0-back: *AUC* = 0.78, *d’* = 1.62; green triangles in Fig. 6B). Together, these results demonstrate that complex visual input interferes with pupillometry-based effort tracking, but crucially also that such interference can be mitigated through dynamic pupil size modeling using Open-DPSM. This should effectively enable unobtrusive, remote, and objective high-temporal resolution tracking of effort with pupil size.

**Figure 6.**
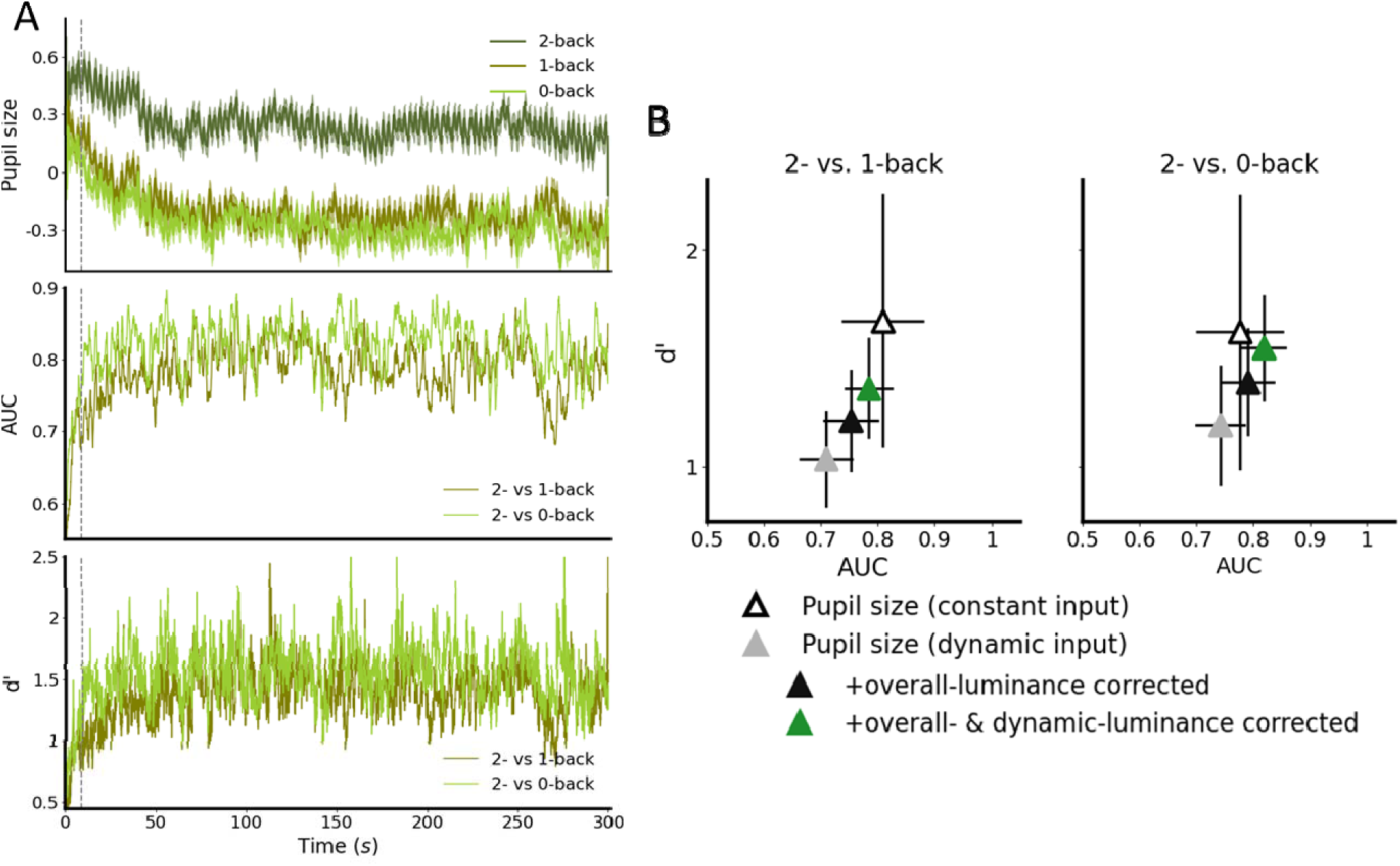
Pupil-based high-temporal resolution effort tracking. *Note: A Example time-series illustrating the univariate discrimination analysis performed at each time point. The top panel shows pupil diameter averaged across n-back conditions. Shaded areas indicate standard error. The middle and bottom panels show the corresponding classification performance (AUC and d0, respectively) for discriminating 2-back from 1-back (olive green) and 2-back from 0-back (light yellow green) at each time point, pooled across participants. The data illustrated here were corrected for luminance with Open-DPSM (corresponding to the green triangles in B; see Supplementary Fig. 1 for the data corrected for overall luminance, data uncorrected for luminance, and data under constant input). Vertical dashed line marks the first 9s, during which cognitive effort was not yet differentiated across n-back conditions; this interval was excluded from all analyses. B Mean classification performance (AUC and d’) averaged across all analyzed time points. Different triangles indicate classification performance of pupil size in dynamic input condition with no luminance correction (grey), corrected with time-averaged luminance model (black), corrected with Open-DPSM (green, which are also the mean of the time-series AUC and d’ in A), as well as classification performance of pupil size in constant input condition (empty triangle). Results for 2- vs. 1-back (left) and 2- vs. 0-back (right) comparisons are shown separately. Error bars indicate the interquartile range (25th–75th percentiles) across all time points.*

## Python package: Dynamic luminance correction with Open-DPSM

The updated Open-DPSM toolbox is available at https://github.com/caiyuqing/Open-DPSM/tree/master/v4. Open-DPSM is implemented in Python (tested with Python 3.12.8) and provides functions for extracting luminance information from dynamic visual input and predicting luminance-driven pupil sizes. The toolbox contains two main classes of functions: luminance extraction and pupil prediction. In addition, a class of function for interactive plotting of results is included. We here provide a summary of the toolbox; a more detailed description can be found on the GitHub page.

### Luminance extraction

The toolbox extracts two types of luminance information from each movie. First, it extracts the overall luminance of each movie, which is later used to estimate time-averaged luminance effects on pupil size. Second, it extracts frame-by-frame luminance changes in a gaze-contingent manner, so that the extracted luminance changes reflect the visual input effectively reaching the retina.

### Pupil prediction

Using overall luminance, the toolbox first predicts *overall* pupil size with a mixed effect model (pupil size ∼ overall luminance + (1| participant)). This step is optional, and users can also provide predicted pupil size obtained from another model of their choice. Then the toolbox predicts visually evoked pupil size *changes* by convolving the extracted luminance changes with a pupil response function. The predicted pupil size from the overall luminance model and the predicted pupil-size changes from the dynamic luminance model are then combined to yield a luminance-driven pupil trace. The model also outputs a luminance-corrected pupil trace, obtained by subtracting the luminance-driven pupil size from the observed pupil size.

### Interactive plotting (optional)

The toolbox can also produce a GUI window to visualize gaze position, extracted luminance changes, observed and predicted pupil size and the luminance-corrected pupil trace over time. Each movie frame can be inspected separately, allowing users to get a better understanding of the data and model results.

## Discussion

The current study evaluated whether and how well pupil size can index cognitive effort under dynamic visual input (i.e., driving movie clips), and whether luminance correction improves its usefulness in such settings. Although luminance is widely recognized as a major confound in pupil-based effort research, few studies have directly evaluated the effectiveness of luminance correction under dynamic visual input, particularly at both aggregate and time-resolved levels, vis-a-vis behavioral and subjective measures. We showed that pupil size increased with task load, in line with behavioural and subjective measures of effort (i.e., worse task performance and higher perceived difficulty). As expected, dynamic visual input reduced the magnitude of effort-related effects relative to constant visual input. However, this reduction could be mitigated by luminance correction, unmasking effort effects on pupil size at both the session level and for moment-to-moment effort tracking over time. Crucially, we here demonstrate that a fully explainable and open-source convolutional model (Open-DPSM) successfully mitigates dynamic changes in visual input, as caused by both display and eye movements during natural viewing. For the aggregate measures (pupil size, accuracy and NASA-TLX score per session), luminance-corrected pupil size numerically slightly outperformed accuracy and NASA-TLX score as measures. More importantly though, combining all three measures (physiology, behavior, subjective experience) yielded clearly the highest dissociation between task loads. Together, we now deem pupillometry-based effort tracking possible across a much wider set of tasks and applications than before. This method is also supported by the updated Open-DPSM toolbox, which is openly available and provides a practical implementation for extracting luminance information, predicting luminance-driven pupil traces, and obtaining luminance-corrected pupil signals in dynamic visual environments.

### 5.1 Dynamic luminance correction allows continuous effort tracking under dynamic visual input

Pupil size is especially suitable for effort tracking, as it can be measured unobtrusively and remotely, and requires no overt behavior from participants. However, pupil size is affected by many factors, most notably luminance changes, which have thus far mostly constrained this power to highly controlled lab environments (Lohani et al., 2019). The current study addresses this constraint by showing that effort-related pupil size change can be recovered under dynamic input using an accessible, open-source, dynamic modeling approach (Open-DPSM), and by directly quantifying its benefit over a simpler overall luminance correction. While we want to reemphasize that this approach generalizes to many more scenarios, driving as an example already shows why this time-resolved tracking is highly important: In driving, effort can fluctuate rapidly with changing traffic complexity, suddenly appearing distractors, and secondary tasks, making moment-to-moment tracking of effort crucial for real-time state monitoring to detect overload/underload states (Engström et al., 2017; Gold et al., 2015; Heikoop et al., 2019; Young & Stanton, 2002), and heightened demands around critical events such as automation handovers (Radhakrishnan et al., 2022) that are safety critical. While previous studies showed that effects of luminance and effort on pupil size could be separated in driving using simplified settings (e.g., with a few different static luminance levels, see Kun et al., 2012; Palinko & Kun, 2011), this has not been translated to unconstrained and dynamic visual input before. The present study directly tested an updated version of Open-DPSM for luminance correction in effort tracking, demonstrating that dynamic luminance modeling can reduce luminance-related influences and recover effort-related pupil responses under dynamic visual input during driving.

Importantly, we here used driving as an instantiation of dynamic and complex visual input, but this method is by no means limited to this example. Any tasks involving dynamic visual input, such as video gaming, education, navigation, sporting, industrial operation, face the same challenge that luminance fluctuations can obscure effort-related pupillary signals unless modeled explicitly (Fanourakis & Chanel, 2022). To facilitate future applications, we provide an updated Open-DPSM that implements luminance correction for pupil signal using only a recording of the visual input together with eye tracking data, offering a practical solution for luminance correction across domains and applications. Very recently, Wang et al. (2026) also extended Open-DPSM framework to emotional video viewing, showing that convolutional-based modeling can capture pupil-size changes driven by both low-level visual features and higher-order emotional factors. Moreover, they also showed that when predicting arousal ratings with pupil size, incorporating the effects of low-level features into the model improved model performance, although performance differed substantially across sessions and participants. Together, these findings suggest that dynamic luminance modeling with Open-DPSM can support pupil-based tracking of different internal states in dynamic visual environments, including both emotional arousal and cognitive effort.

### 5.2 Cognitive effort is best captured according to its definition: as a multifaceted construct

Effort is defined multidimensionally with distinct and complementary physiological, behavioural, and subjective effects. When measuring effort in applied settings, the common trend is to adopt multiple measurements (e.g., Armougum et al., 2020; Johannessen et al., 2020). Our data support this view, as the effort-inducing task load could best be separated by combining measures tapping into each dimension. Physiological (pupil size), behavioural (accuracy) and subjective (NASA-TLX) indices were linked, but only moderately, indicating that they all capture unique variance. Accordingly, previous studies have shown that partial dissociations among measurement dimensions may reflect different outcomes of the same underlying cognitive demands. For instance, perceived workload does not necessarily translate into behavioral costs (Longo et al., 2022) and physiological measures can remain sensitive at behavioural ceiling (McLaughlin & Van Engen, 2020). On the other hand, adding additional physiological channels (e.g., combining EEG and pupil size) does not always improve effort prediction (Aygun et al., 2022; Borys et al., 2017; but see Khedher, 2019; Rozado & Dünser, 2015). Taken together, these findings suggest that multimodal assessment combining all three modalities is ideal for an optimal representation of effort when feasible.

However, in many applied settings, assessing all dimensions is impractical. We showed that luminance-corrected pupil size provided slightly better classification performance than accuracy and NASA-TLX. Previous studies also reported pupil size to outperform other physiological signals (Aygun, Lyu, et al., 2022; Aygun, Nguyen, et al., 2022; Schiff & Foa, 1875), indicating that it can be a reliable standalone indicator with proper luminance correction. Nevertheless, measure selection should depend on the question asked: obviously, behavioural measures are most appropriate when similar behaviour is the primary outcome, self-reports are preferred when subjective experience is central, and physiological signals are advantageous when continuous, unobtrusive tracking is required. Among physiological measures, pupillometry comes at a relatively low cost, and is therefore practical in the applied and fundamental contexts alike.

### 5.3 Limitations and future directions

The present study used driving movies to capture the real-world visual complexity of driving, but this setup remains distinct from real driving, where measurement of pupil size can additionally be affected by additional factors, such as head and body movements, which affect pupil size as well or pose challenges for valid data acquisition. As we excluded sessions with low quality (more than 40% missing data) for this proof-of-concept, our conclusions are based on rather clean data. Future work should therefore test the proposed luminance correction under even noisier conditions. In principle, this method should allow the capture of moment-to-moment fluctuations in effort and thus under-/overload and therefore also predict behavior, such as overlooking a critical road sign, not responding to a critical alarm in industrial operations, etc. This of course remains to be tested. Finally, although luminance changes are by far the most important drivers of pupil size change, additional factors beyond effort are reflected in the pupil (Strauch et al., 2022). For instance, other low-level visual features, such as spatial frequency, colour and orientation can also influence pupil size (Barbur et al., 1992; Hu et al., 2019; Young et al., 1995) and the pupil responds to changing depths as well (Pielage et al., 2022). In addition, higher-level time-varying annotations, such as continuous arousal ratings, can be incorporated into the convolutional model framework as additional input traces, as recently demonstrated by Wang et al. (2026). Depending on the research question, this may also allow researchers to model psychological factors of interest or account for additional psychological influences when isolating a targeted higher-order pupil component. Other confounds include measurement related issues, such as the pupil foreshortening error (Gagl et al., 2011; Hayes & Petrov, 2016; Stribos et al., in press), should also be accounted for. As mentioned in Cai et al. (2024), Open-DPSM is fully open and in principle able to incorporate other factors affecting pupil size (through effort or not), which we see as an important future direction for further improving pupil-based effort estimation in dynamic, real-world settings.

## Conclusion

The current study illustrated that pupil size well indexes cognitive effort under dynamic visual input when (dynamic) luminance correction is applied. Luminance effects have long been considered the central roadblock to applying and leveraging pupil size change as a marker of effort beyond the highly controlled laboratory setting in which effort-related effects on the pupil were established. We validated a method that overcomes this roadblock, not only on the aggregate level (i.e., average pupil size per session) but also over time (from time point to time point). Compared with behavioral and subjective indices, luminance-corrected pupil size provided a sensitive standalone measurement. In addition, combining physiological, behavioral, and subjective measures yielded the strongest classification performance, indicating that effort is multifaceted and can be best measured by a combination of multimodal indicators, including pupil size.

## Open Practices Statement

All materials, data, analysis scripts and supplementary materials are available at https://osf.io/2tbpk and the experiment was not preregistered.

## Acknowledgement

Yuqing Cai was supported by a China Scholarship Council (CSC) scholarship. Christoph Strauch received funding from the Dutch Research Council (NWO), Veni grant (VI.Veni.241G.005, 10.61686/EEGGV23807).

