## Supplementary materials for "Luminance-corrected Pupillometry Reliably Tracks Effort in Dynamic Driving Environments"

**Supplementary Table 1.**

Repeated-measures ANOVA results for accuracy and NASA-TLX

| Dependent variable | Effect | df1 | df2 | *F* | *p* | *ηp²* |
|---|---|---|---|---|---|---|
| Accuracy | N-back (0-, 1-, 2-back) | 2 | 44 | 31.26 | < 0.001 | 0.35 |
| | Visual input (constant, dynamic) | 1 | 22 | 3.51 | 0.07 | 0.01 |
| | Interaction | 2 | 44 | 0.17 | 0.84 | 0.001 |
| NASA-TLX | N-back (0-, 1-, 2-back) | 2 | 44 | 81.22 | < 0.001 | 0.36 |
| | Visual input (constant, dynamic) | 1 | 22 | 0.35 | 0.56 | 0.002 |
| | Interaction | 2 | 44 | 0.41 | 0.67 | 0.004 |

*Note: p-values two-tailed; ηp² = partial eta squared.*

**Supplementary Table 2.**

Post-hoc pairwise comparisons across n-back levels (within each visual input condition)

A. Accuracy

| Visual input | Comparison | df | *t* | *p* | *d* | *95% CI* |
|---|---|---|---|---|---|---|
| Dynamic | 2 vs 1 | 22 | 4.65 | < 0.001 | 0.97 | [0.04, 0.11] |
| Dynamic | 2 vs 0 | 22 | 5.63 | < 0.001 | 1.17 | [0.06, 0.12] |
| Dynamic | 1 vs 0 | 22 | 1.02 | 0.32 | 0.21 | [-0.01,0.03] |
| Constant | 2 vs 1 | 22 | 4.07 | 0.001 | 0.85 | [0.04, 0.13] |
| Constant | 2 vs 0 | 22 | 5.22 | < 0.001 | 1.09 | [0.06, 0.14] |
| Constant | 1 vs 0 | 22 | 1.74 | 0.10 | 0.36, | [-0.002, 0.02] |

B. NASA-TLX

| Visual input | Comparison | df | *t* | *p* | *d* | *95% CI* |
|---|---|---|---|---|---|---|
| Dynamic | 2 vs 1 | 22 | -8.68 | < 0.001 | -1.81 | [-4.28, -2.63] |
| Dynamic | 2 vs 0 | 22 | -8.25 | < 0.001 | -1.72 | [-5.91, -3.53] |
| Dynamic | 1 vs 0 | 22 | -3.15 | 0.005 | -0.66 | [-2.10, -0.43] |
| Constant | 2 vs 1 | 22 | -5.27 | < 0.001 | -1.10 | [-5.04, -2.19] |
| Constant | 2 vs 0 | 22 | -6.44 | < .001 | -1.34 | [-5.45, -2.79] |
| Constant | 1 vs 0 | 22 | -0.76 | 0.46 | -0.16 | [-1.89, 0.88] |

*Note: p-values two-tailed; d = Cohen's d*

**Supplementary Table 3.**

Repeated-measures ANOVA results for mean pupil size

| Condition | Effect | df1 | df2 | *F* | *p* | *ηp²* |
|---|---|---|---|---|---|---|
| Constant input | N-back (0-, 1-, 2-back) | 2 | 46 | 19.98 | < 0.001 | 0.46 |
| Dynamic input, uncorrected | N-back (0-, 1-, 2-back) | 2 | 62 | 23.03 | < 0.001 | 0.42 |
| Dynamic input, overall luminance-corrected | N-back (0-, 1-, 2-back) | 2 | 62 | 47.21 | < 0.001 | 0.59 |

*Note: p-values two-tailed; ηp² = partial eta squared.*

**Supplementary Table 4.**

Post-hoc pairwise comparisons across n-back levels for mean pupil size

| Condition | Comparison | df | *t* | *p* | *d* | *95% CI* |
|---|---|---|---|---|---|---|
| Constant input | 2 vs 1 | 23 | -5.25 | < 0.001 | -1.07 | [-0.82, -0.36] |
| | 2 vs 0 | 23 | -5.36 | < 0.001 | -1.09 | [-0.73, -0.32] |
| | 1 vs 0 | 23 | 0.64 | 0.53 | 0.13 | [-0.14, 0.26] |
| Dynamic input, uncorrected | 2 vs 1 | 31 | -5.24 | < 0.001 | -0.93 | [-0.58, -0.26] |
| | 2 vs 0 | 31 | -5.54 | < 0.001 | -0.98 | [-0.68, -0.31] |
| | 1 vs 0 | 31 | -1.14 | 0.26 | -0.2 | [-0.20, 0.06] |
| Dynamic input, over all luminance–corrected | 2 vs 1 | 31 | -7.78 | < 0.001 | -1.37 | [-0.55, -0.32] |
| | 2 vs 0 | 31 | -7.82 | < 0.001 | -1.38 | [-0.63, -0.37] |
| | 1 vs 0 | 31 | -1.32 | 0.33 | -0.23 | [-0.16, 0.03] |

*Note: p-values two-tailed; d = Cohen's d*

**Supplementary Table 5.**

Discriminability of 2-back vs. 1-/0-back conditions (AUC and d')

| Visual input | Measure | Comparisons | | | |
|---|---|---|---|---|---|
| | | 2 vs 1 | | 2 vs 0 | |
| | | AUC | d' | AUC | d' |
| Constant | Accuracy | 0.84 | 1.61 | 0.89 | 1.99 |
| Constant | NASA-TLX | 0.79 | 1.49 | 0.87 | 2.11 |
| Constant | Pupil size | 0.95 | 2.51 | 0.90 | 2.31 |
| Constant | Accuracy + NASATLX + Pupil size | 0.98 | 3.03 | 0.95 | 5.35 |
| Dynamic | Accuracy | 0.77 | 1.22 | 0.84 | 1.70 |
| Dynamic | NASA-TLX | 0.81 | 1.43 | 0.88 | 1.83 |
| Dynamic | Pupil size | 0.80 | 1.31 | 0.81 | 1.90 |
| Dynamic | Pupil size (overall luminance–corrected) | 0.89 | 1.94 | 0.92 | 1.99 |
| Dynamic | Accuracy + NASATLX + Pupil size (overall luminance–corrected) | 0.92 | 2.02 | 0.98 | 2.94 |

**Supplementary Figure 1.**

Discrimination analysis performed at each time point

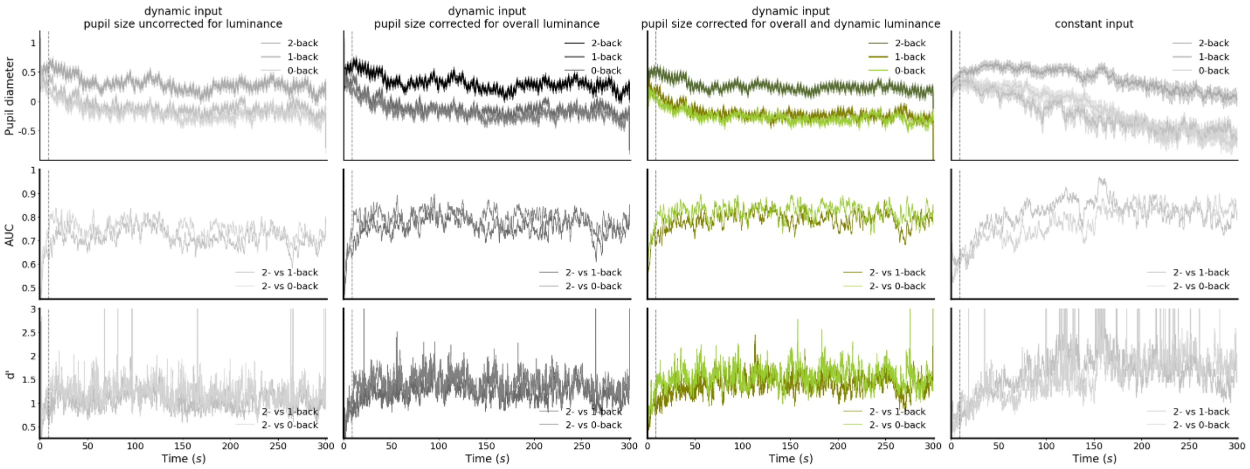

*Note: same as Fig. 6A but with different pupil data. From left to right: data uncorrected for luminance, data corrected for overall luminance, data corrected for overall and dynamic luminance (with Open-DPSM), and data under constant input.*
